# Analyzing the effects of altitudes on the metabolic diversity and forskolin content in *Coleus forskohlii* roots by HPTLC and HPLC

**DOI:** 10.1101/2020.04.15.041970

**Authors:** Pawan Singh Rana, Pooja Saklani, Chandresh Chandel

## Abstract

**Introduction:** *Coleus forskohlii* is an important medicinal plant native to India. It grows wild in a wide range of altitude in the Indian Himalayan region and contains some important phytochcemicals which possess remarkable medicinal properties. The plant contains terpenoid like Forskolin in its roots.

**Objective:** Considering the medicinal importance of *C. forskohlii*, being the only source of Forskolinand its availability over a wide altitude range, the effect of altitude on the metabolic diversity and forskolin content was assessed using the HPTLC and HPLC.

**Methods:** Five populations of *Coleus forskohlii* collected from five locations of varying altitude from Uttarakhand, India. The plant roots were extracted with methanol by soxhlet extraction. The metabolic diversity was analyzed by employing HPTLC fingerprinting while forskolin was extracted and quantified by HPLC.

**Results:** Significant differences were observed in the phytochemical composition through the HPTLC chromatograms among the studied population across the altitudes. The Gopeshwar population (1488m) showed highest number of bands on HPTLC chromatogram each of which corresponds to acompound. Results of HPLC shows differences in forskolin quantity in studied populations and the Piaplkoti population(1339m) showed highest forskolin accumulation.

**Conclusion:** The present study confirms that altitude and changing environment affects the nature and quantity of secondary metabolites in *C. forskohlii* and the environmental conditions might be instrumental factor for intraspecific metabolic diversity. The Pipalkoti (1339m) and Gopeshwar (1488m) populations were found suitable for the forskolin production as well as other metabolites and these two populations can be propagated for commercial use.

## 1. Introduction

*Coleus forskohlii* commonly known as Patharchur, is a perennial herb grows in subtropical conditions. The plant is endemic to India grows in almost all the south east Asian countries like Sri Lanka, Nepal, Thailand, Pakistan and Bhutan**(Valdes et al., 1987)**. It is well documented for its use to cure gastric problems, inflammation, glaucoma, hypertension and piles (**Srivastava et al., 2002; Shah et al., 1980; De Souza et al., 1986**). The plant grows wild in Uttarakhand, and being used by locals and tribal population for curing the external wounds, ulcer, cough, throat infections and psoriasis (**Kotia et al., 2014**). Although the whole plant is medicinally potent but the roots have greater value as they have been reported to accumulate an important diterpenoid Forskolin. Forskolin possess a unique property of activating the adenylate cyclase which is regulator of cyclic AMP levels in the cell (**Seamon et al., 1981**). The activation of cyclic AMP has great biochemical and physiological effects. The plant is also well known for its antioxidant and antiageing potential **(Adachi et al., 1996)**. Limited studies on the HPTLC analysis of *Coleus forskohlii* have been reported from different altitudes (**Srivastava et al., 2017**). Although work have been done to estimate the forskolin quantity and the chemotypic variability of forskolin in the cultivated and wild populations of *Coleus forskohlii* (**Tamboli et al., 2013; Ahmad et al., 2011**), but efforts have not been made to analyze the total metabolic diversity. Considering its medicinal importance, it had been studied time to time and region to region by various workers but the chemotypic variability by HPTLC in the Garhwal region of Uttarakhand (India) had not yet been well documented. As the plant grows along a wide range of altitudes, five populations of *Coleus forskohlii* were collected from five different locations of different altitudes. Considering the medicinal importance of forskolin, all the five populations were also analyzed for forskolin content using HPLC. Therefore, the present study was designed to analyze the level of metabolic diversity by HPTLC and forskolin content by HPLC in the five populations of *Coleus forskohlii* from different locations of Uttarakhand, India.

## 2. Materials and Methods

### 2.1 Plant material

Whole plant samples were collected from all the selected five locations. The numbers of samples were limited to 6-10 from each of the location keeping in mind the sustainable approach. The plants were thoroughly washed under running tap water and roots were detached. These, roots were further dried at the room temperature and were powdered by using liquid nitrogen in mortar pestle. Botanical identity of the sample was authenticated by Botanical Survey of India (Dehradun), Accession No. 118603.

### 2.2 Chemicals and Reagents

All the chemicals used were of analytical grade. The chromatographic plates used were aluminium backed silica gel plate 60F254 HPTLC plate (E. Merck Ltd, Darmstadt, Germany). Sample applicator CAMAG Automatic TLC Sampler 4 (ATS 4), twin trough chamber (20×10cm) for development, Whatmann filter paper (No 41), CAMAG scanner and UV chamber for visualization (Short and long wavelength), dipping tank and plates dryer were used. For HPLC, HiQ Sil C18-HS column was used, with Systronics HPLC system fitted with a photo diode array UV detector.

### 2.3 Extraction of plant material

The coarsely powdered roots (50gm) were extracted with 300ml of methanol in a Soxhlet extractor for about 6 hr. The extracts so obtained were further concentrated by evaporating the solvent on a water bath. The solvent free extract so obtained was further dried using a lyophilizer. The dried extracts than dissolved in HPLC grade methanol and stored sealed bottles for further analysis.

### 2.4 Phytochemical Screening

The concentrated extracts were subjected to chemical test as per **Devi et al., 2014; Kumar et al., 2013**.

#### 2.4.1 Selection of solvent system

Solvent system for HPTLC was selected using many mobile solvent combinations **(Wagner & Bladt, 1996)** and finally Dichloromethane: Methanol: Water (7:3:0.3) was selected.

#### 2.4.2 HPTLC fingerprinting profile

HPTLC fingerprints of each of the five populations were developed using the CAMAG HPTLC system.

##### 2.4.2.1 Sample preparation

The extracts were made at a concentration of 5mg ml^-1^ in HPLC methanol and filtered with Whatman filter No. 41. Further, extracts were diluted 10 fold with the solvent again.

##### 2.4.2.2 Sample application

The samples were band-wise applied on the HPTLC plates using the CAMAG ATS-4 applicator. The samples were applied in 15 tracks (band length 8mm) at a distance of 8mm from the lower edge and 20mm from left edge with an application volume of 1µl, 2µl, 4µl for each extract.

##### 2.4.2.3 Chromatography

Extracts of the each of five populations of *Coleus forskohlii* were separated in a Twin Trough Chamber (20×10cm) with Dichloromethane: Methane: Water (7:3:0.3) up to a migration distance of 80mm. Further, the plates were dried for 5min at room temperature. The chromatograms were neutralized with potassium hydroxide for 4 h followed by cold air stream for 20 min.

##### 2.4.2.5 Post chromatographic Derivatization

The plates were dipped in Anisaldehyde Sulphuric Acid Reagent (ASR) using Chromatogram immersion device at an immersion speed of 5cm/s for 1s. Further, the plate was heated at 100°C for 3 min after using a tissue heater.

##### 2.4.2.6 Documentation

The finally developed plate was observed under HPTLC visualizer at 254nm, 366nm (both before derivatization), 366nm and white light (540nm) after derivatization.

### 2.5 Forskolin Extraction and confirmation by TLC

Forskolin was extracted by the method adopted from **Kumar & Spandana, 2013**. A reddish brown colored powder obtained. Presence of forskolin was further confirmed by performing thin layer chromatography with the standard.

#### 2.5.1 Forskolin estimation by HPLC

The stock solution of concentration 10 µg/ml was prepared by dissolving 10µg forskolin in 0.5ml HPLC-grade methanol followed by sonication for 10 minutes and the resulting volume was made up to 1ml with the methanol. The same method was followed to prepare the sample solutions.10µl of all the samples were mixed in 1 ml of methanol and then vortex. They were then brought to filter through 0.2µm of filters. The standard and sample solutions were filtered through 0.22μm PVDF-syringe filter and the mobile phase was degassed before the injection of the solutions. The mobile phase contains Methanol: Acetonitrile: Water in the ratio of 60:30:10 the flow rate was adjusted to 1.0 ml/min, the column was thermostatically controlled at 25°C and the injection volume was kept at 20 μl and isocratic elution was performed. Total analysis time per sample was 10 min. HPLC chromatograms were detected using a photo diode array UV detector at single wavelengths of 220 nm according to absorption maxima of analysed compounds. Each compound was identified by its retention time and by spiking with standards under the same conditions. The quantification of the sample was done by the measurement of the integrated peak.

## 3. Results

The samples of *Coleus forskohlii* were collected from five different locations of Garhwal region located between 600-2000m altitudes (Table 1). No morphological variations were observed visually. In the earlier studies methanol was reported as best solvents for extraction and various activities hence methanol was chosen for extracting the metabolites. The phytochemical screening revealed presence of alkaloids, flavonoids, phenols, terpenoids, tannins, coumarins, quinones and proteins while steroids, xanthoproteins, glycosides, saponins were absent (Table 2) in all the five population of *C. forskohlii*.

**Table 1.**
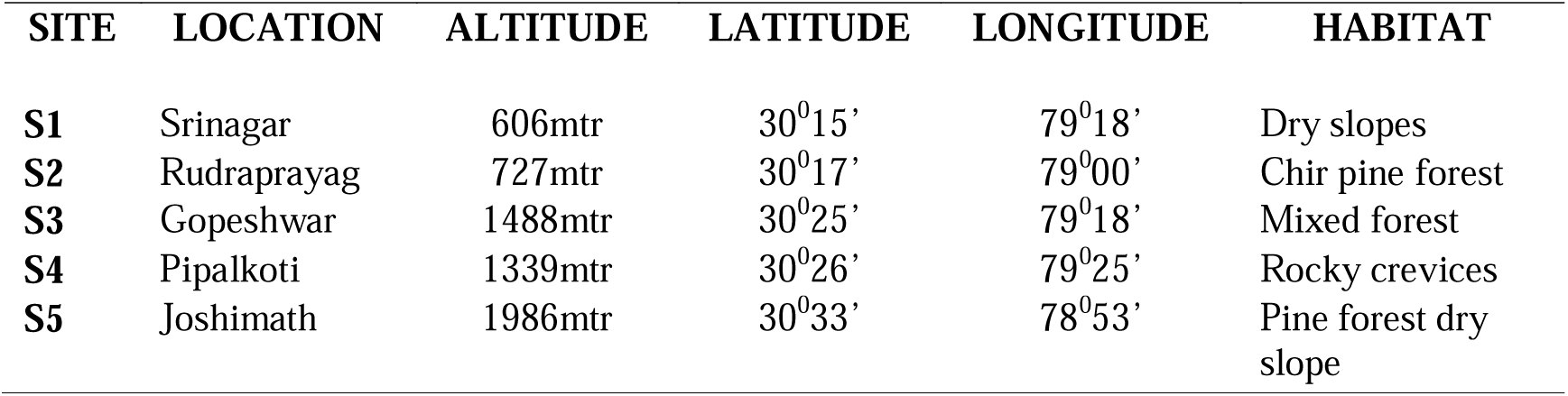
Details of sampling sites.

**Table 2.** Phytochemical screening of methanolic extracts of *C. forskohlii* roots.

| S. No. | Tests | Result |
| --- | --- | --- |
| 1 | **Alkaloids** | +ve |
| 2 | **Flavonoids** | +ve |
| 3 | **Phenols** | +ve |
| 4 | **Terpenoids** | +ve |
| 5 | **Steroids** | -ve |
| 6 | **Coumarins** | +ve |
| 7 | **Tannins** | +ve |
| 8 | **Xanthoproteins** | -ve |
| 9 | **Quinones** | +ve |
| 10 | **Carbohydrates** | -ve |
| 11 | **Glycosides** | -ve |
| 12 | **Proteins** | +ve |
| 13 | **Saponins** | -ve |

### 3.1 TLC analysis

The TLC analysis of all the five population showed highest number of band in the combination of Dichloromethane: Methanol: Water (7: 3: 0.3), hence this combination was used for HPTLC fingerprinting.

### 3.2 HPTLC fingerprinting analysis

The chromatogram of HPTLC finger printing revealed the presence of various metabolites when viewed under UV light and white light (Fig. 1 and 2). The HPTLC fingerprints showed highest number of peaks in the populations of higher altitudes, 8 and 5 peaks respectively were recorded in the CFr04 (Gopeshwar population) and CFr05 (Joshimath population), when observed under 366nm while 6 and 4 peaks were observed for both of these populations respectively at 540nm (Table 3). It was quite clear from the HPTLC fingerprints that the populations of lower altitude Srinagar and Rudraprayag (CFr01 and CFr02) had lesser band (3 and 4 respectively at 366nm and 4 at 540nm) as compare to those of a higher altitude.

**Table 3.**
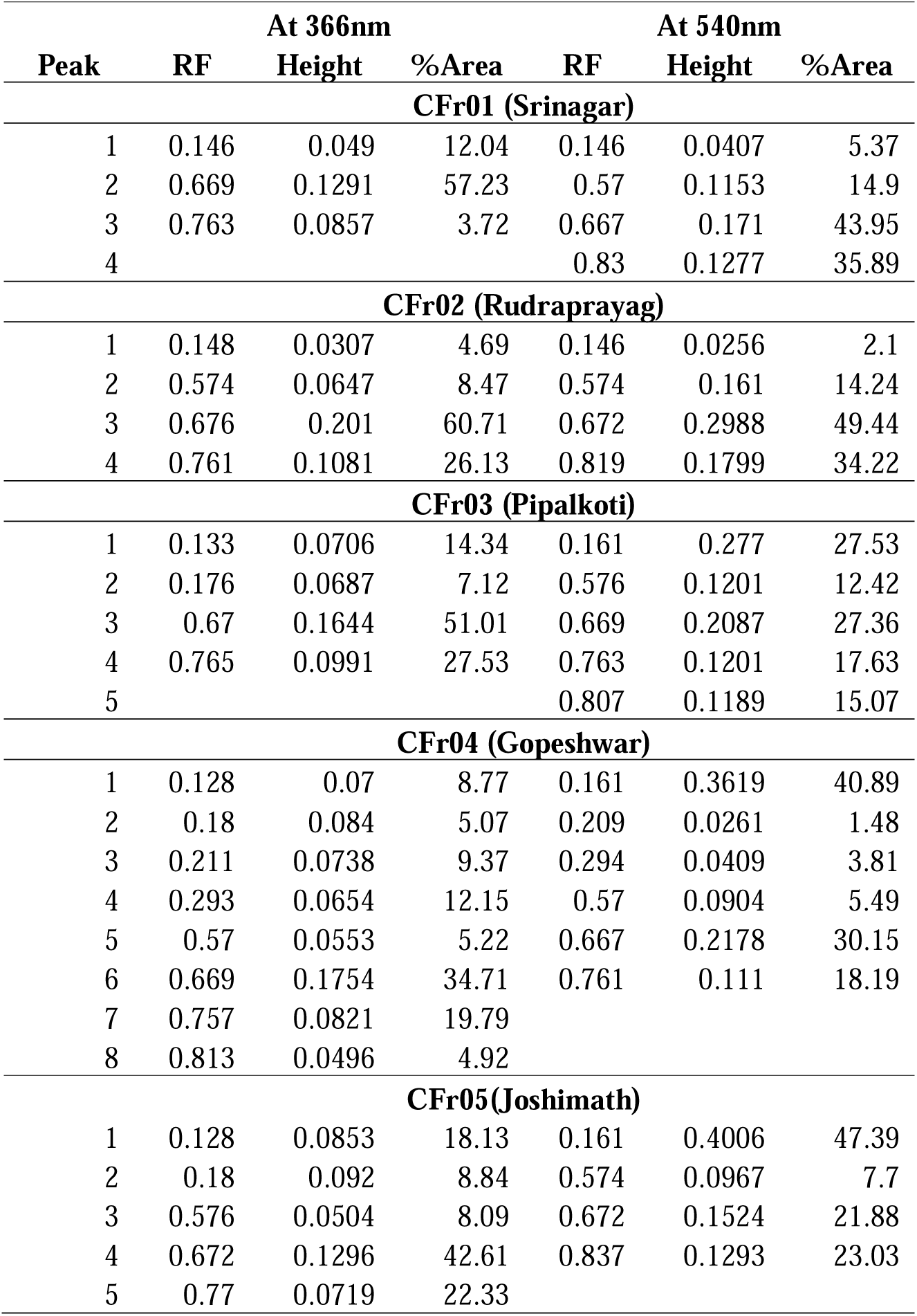
Peaks for phytochemicals observed at 366nm and 540nm.

**Fig. 1.**
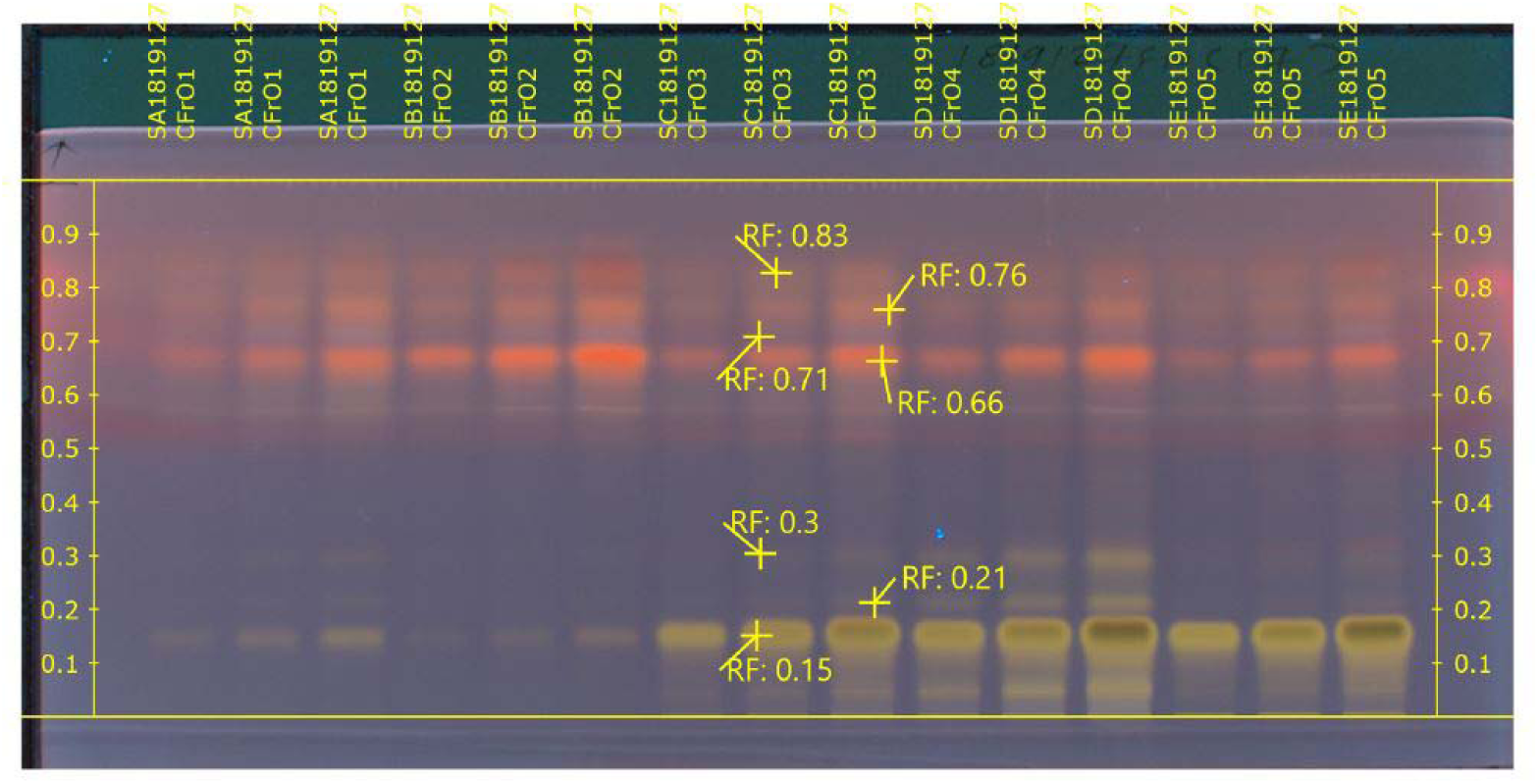
Chromatogram showing separated phytochemicals when viewed under 366nm.

**Fig. 2.**
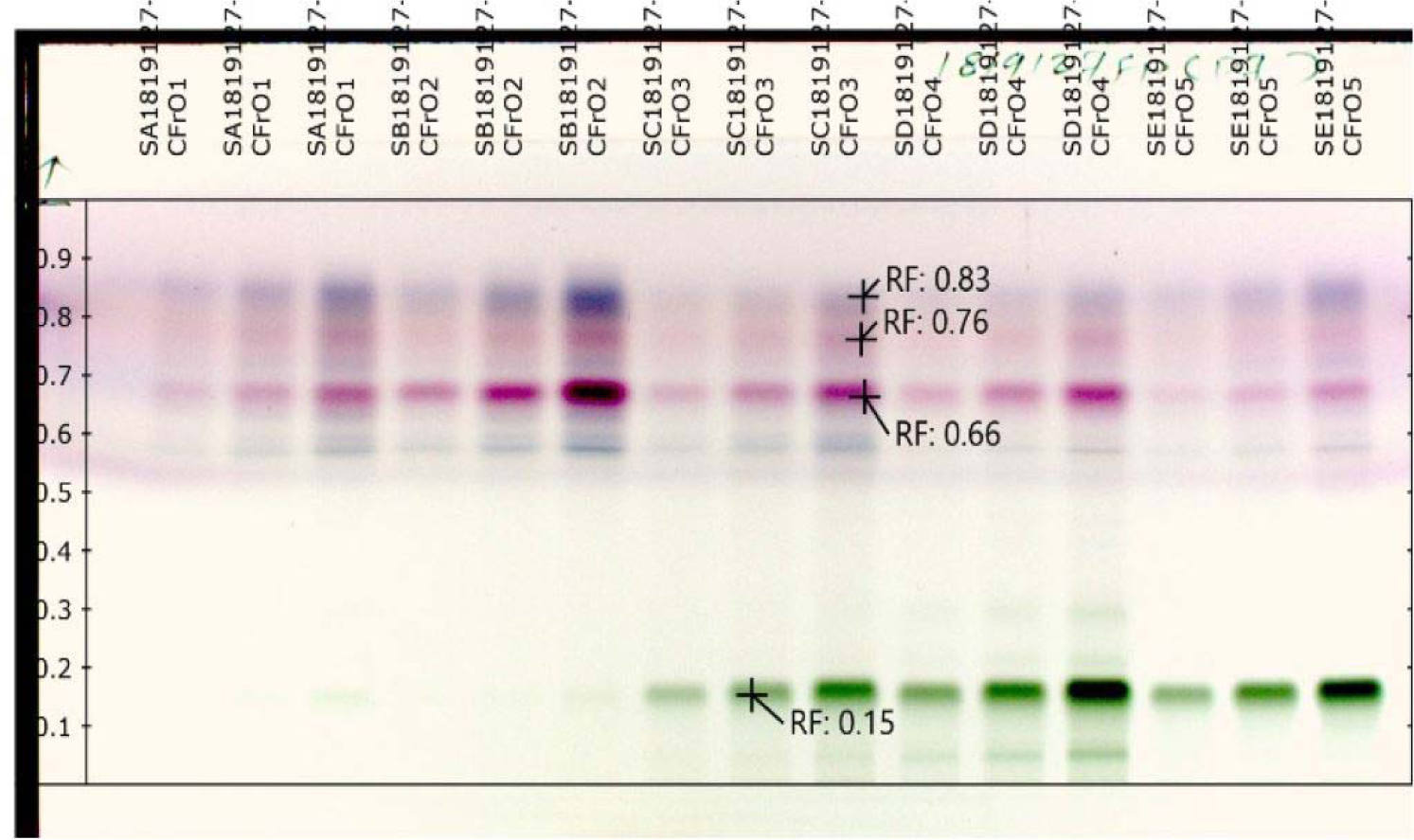
Chromatogram showing separated phytochemicals when observed under 540nm.

**Fig. 3.**
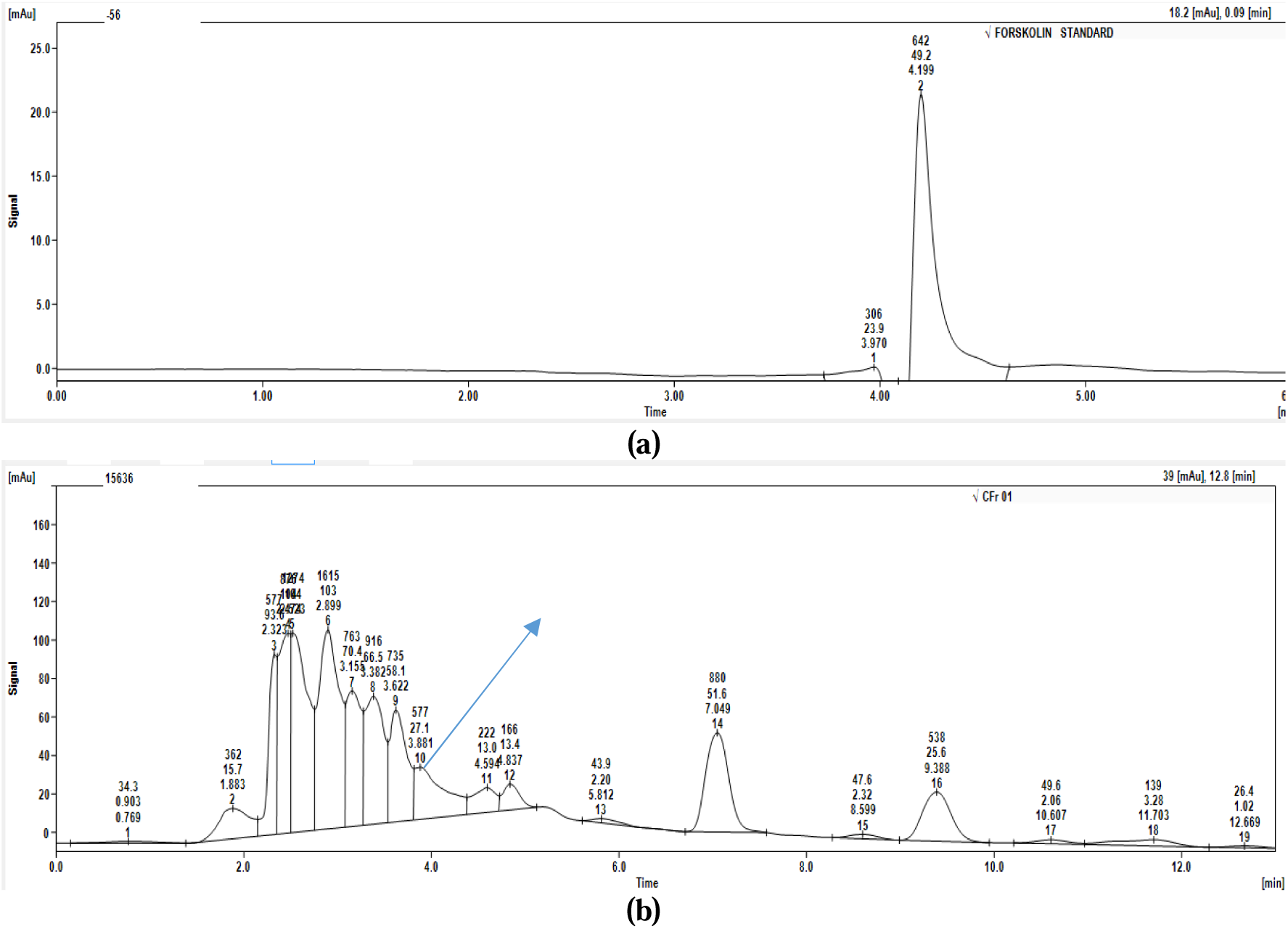
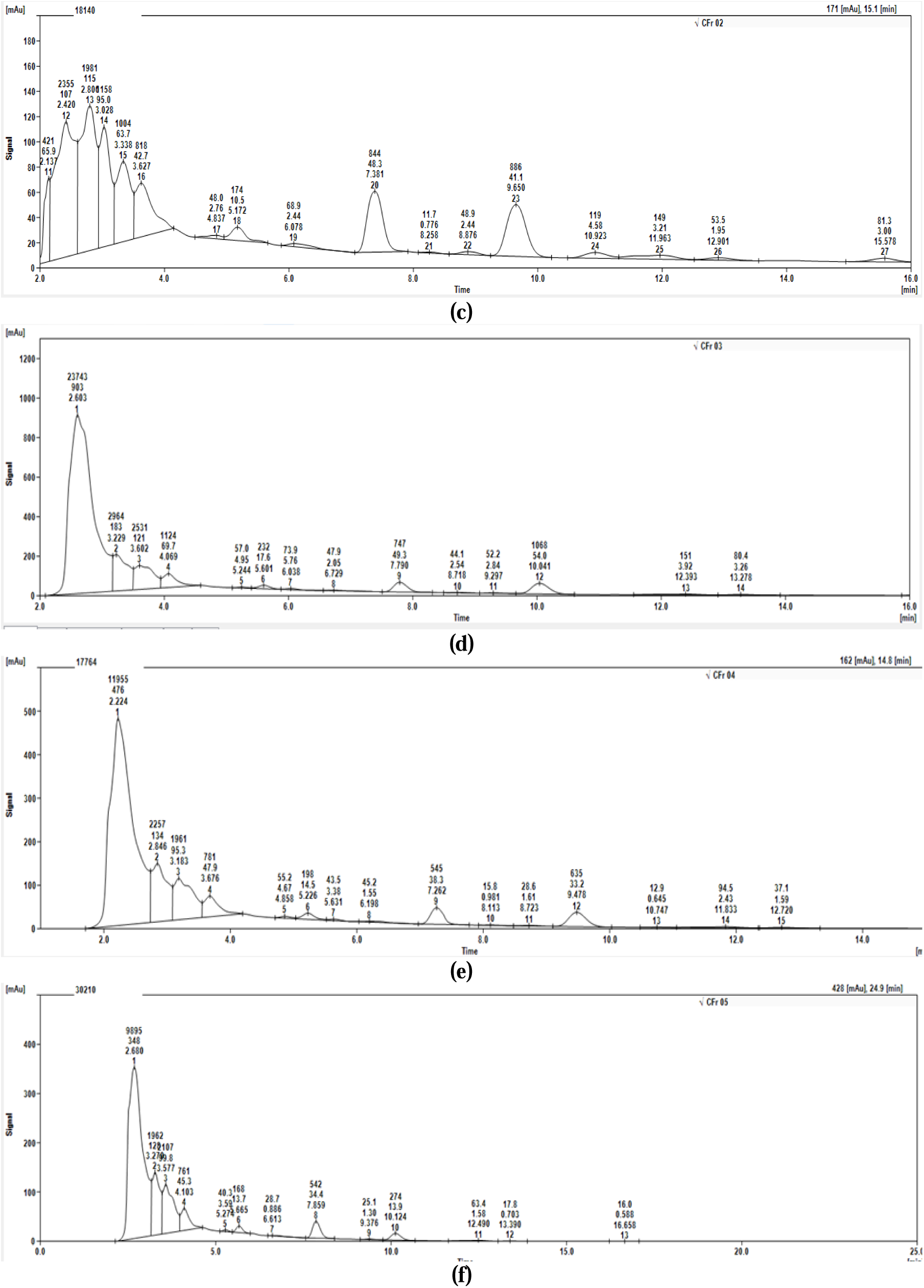
HPLC chromatograms (a) Forskolin standard (b) CFr01, (c) CFr02, (d)CFr03, (e) CFr04, (f) CFr05.

Table 4 shows comparison of the different RF values that has been detected in the chromatogram, it is quite evident that there are differences in the RF values (each referring to a compound) among the five populations. The compound at an RF 0.14, 0.57 and 0.67 were observed in all the populations while those at 0.21 and 0.29were found only in the CFr04 (Gopeshwar population). Out of the total 9 compounds observed, 8 were reported in Gopeshwar population (1488m altitude) and 6 in the Joshimath population (1986m altitude).

**Table 4.** Showing the presence and absence of compounds (RF) in all the populations of *C. forksolhii*.

| Compound (RF) | CFr01 | CFr02 | CFr03 | CFr04 | CFr05 |
| --- | --- | --- | --- | --- | --- |
| 0.14 | + | + | + | + | + |
| 0.18 | -- | -- | + | + | + |
| 0.21 | -- | -- | -- | + | -- |
| 0.29 | -- | -- | -- | + | -- |
| 0.57 | + | + | + | + | + |
| 0.67 | + | + | + | + | + |
| 0.76 | -- | + | + | + | + |
| 0.81 | -- | -- | + | + | -- |
| 0.83 | + | + | -- | -- | + |

### 3.3 Forskolin estimation

Extracted forskolin powder was dissolved in HPLC grade methanol and its presence was confirmed by thin layer chromatography where a compact pink color band was obtained at an RF of 0.25 as that of the standard. HPLC analysis shows presence of forskolin in all the samples. The Forskolin content was calculated by comparing the peak area of standard and test samples. Highest forskolin content was observed in the Pipalkoti population, followed by Rudraprayag, Gopeshwar and Joshimath while lowest value was recorded in the Srinagar population (Table 5).

**Table 5.** Forskolin content in the five populations.

| Sample Code | Location | Altitude | Peak Area | Quantity<br>($\mu\text{g gm}^{-1}\text{DW}$) |
| --- | --- | --- | --- | --- |
| CFr01 | Srinagar | 606m | 576,802 | 8.9 |
| CFr02 | Rudraprayag | 727m | 818,476 | 12.74 |
| CFr03 | Pipalkoti | 1339m | 1123,611 | 17.49 |
| CFr04 | Gopeshwar | 1488m | 781,270 | 12.16 |
| CFr05 | Joshimath | 1986m | 760,908 | 11.84 |

## 4. Discussion

It was evident from the results that the compounds present in the populations of lower altitude were also present in the higher altitude populations, while some of those which are reported in the higher altitude populations were absent in lower altitude populations. The results suggest towards a high intraspecific metabolic diversity in the roots of *Coleus forskohlii* among the studied populations. The different RF values in the HPTLC chromatogram clearly indicate the differences in metabolites and their quantities. This may be attributed to the microclimate of the region, genetic character of the plant, altitude of the plant origin, light intensity and growing season of the plant. The effect of geographical conditions on the phytochemicals have already been studied in many plants such as *Zinzibar officinale* **(Ghasemzadeh et al., 2010)**, *Valeriana jatamansi* **(Jugran et al., 2014)**, *Hedychium spicatum* **(Rawat et al., 2011)** and *Crocus sativus* **(Mangal et al., 2018).** With the change in altitude the temperature changes and it may affect the phytochemical composition of the plant, effect of temperature on the phytochemical contents in the *Azadirachta indica* has already been studied **(Vats, 2015)**. The HPTLC fingerprinting results showed metabolic variability as some of the spots were not detected in all the samples, similar results have been observed in *Stevia rebaudiana* (**Chester et al., 2012**) while **Tamboli et al., 2013**have reported negligible metabolic diversit. Forskolin content in all the five populations was estimated by HPLC showed a significant difference (<0.05) across the altitudes. Highest forskolin content was reported in the CFr03 (Pipalkoti population) indicates that it is the best location for forskolin production. The difference in the forskolin content from different ecogeographical regions have already been reported form different parts of India **(Ahmad et al., 2011; Narayanan et al., 2002; Saleem et al., 2005; Schaneberg et al., 2003)**.

## 5. Conclusion

The present study reveals the metabolic diversity of *Coleus forskohlii* growing at different altitudes. It suggested that there is a significant difference in the secondary metabolite profile of the plants growing at different altitude and habitat. The forskolin quantity also varies with the altitudes and habitat which further signifies that the microclimate and altitudes may have some role in the production of secondary metabolites of *Coleus forskohlii* including forskolin. Here, the study reports Pipalkoti location (1339m) as best for forskolin production while highest metabolic diversity was observed at Gopeshwar location (1488m). The molecular studies could be carried out to confirm the genetic basis of variability. Hence, these locations can be explored for the mass cultivation and these two populations can be recommended for commercial cultivation of *Coleus forskohlii* in Uttarakhand, India.

## Acknowledgement

We gratefully acknowledge the Uttarakhand Council for Biotechnology for funding through UCB/R&D Project/2017/47.

